# Chronic ethanol exposure produces long-lasting, subregion-specific physiological adaptations in RMTg-projecting mPFC neurons

**DOI:** 10.1101/2024.05.06.592759

**Authors:** Kathryn R Przybysz, Joel E Shillinglaw, Shannon R Wheeler, Elizabeth J Glover

## Abstract

Chronic ethanol exposure produces neuroadaptations in the medial prefrontal cortex (mPFC) which facilitate the maladaptive behaviors interfering with recovery from alcohol use disorder. Despite evidence that different cortico-subcortical projections play distinct roles in behavior, few studies have examined the physiological effects of chronic ethanol at the circuit level. The rostromedial tegmental nucleus (RMTg) is a GABAergic midbrain region involved in aversive signaling and is functionally altered by chronic ethanol exposure. Our recent work identified a dense input from the mPFC to the RMTg, yet the effects of chronic ethanol exposure on this circuitry is unknown. In the current study, we examined physiological changes after chronic ethanol exposure in prelimbic (PL) and infralimbic (IL) mPFC neurons projecting to the RMTg. Adult male Long-Evans rats were injected with fluorescent retrobeads into the RMTg and rendered dependent using a 14-day chronic intermittent ethanol (CIE) vapor exposure paradigm. Whole-cell patch-clamp electrophysiological recordings were performed in fluorescently-labeled (RMTg-projecting) and -unlabeled (projection-undefined) layer 5 pyramidal neurons 7-10 days following ethanol exposure. CIE significantly increased intrinsic excitability as well as excitatory and inhibitory synaptic drive in RMTg-projecting IL neurons. In contrast, no lasting changes in excitability were observed in RMTg-projecting PL neurons, although a CIE-induced reduction in excitability was observed in projection-undefined PL neurons. CIE also increased excitatory synaptic drive in RMTg-projecting PL neurons. These data uncover novel subregion- and circuit-specific neuroadaptations in the mPFC following chronic ethanol exposure and reveal that the IL mPFC-RMTg projection is uniquely vulnerable to long-lasting effects of chronic ethanol.

## Introduction

Alcohol use disorder (AUD) is a chronic, relapsing disorder characterized by loss of control over alcohol consumption, a negative affective state in the absence of alcohol (i.e., withdrawal), and deficits in decision-making related to seeking and using alcohol which frequently occur despite negative consequences. It has been proposed that, in an alcohol-dependent state, a motivational switch occurs during which the decision to drink alcohol is driven by relief from the aversive effects of withdrawal, rather than the rewarding effects of intoxication (see Koob, 2021; Sinha, 2013 for review). Each of these cognitive and behavioral facets of AUD is reliant, at least in part, on neuroadaptations in the medial prefrontal cortex (mPFC) that occur in response chronic exposure to alcohol (Cyders et al., 2014; Jasinska et al., 2014; Koob & Volkow, 2016; Le Berre et al., 2017; Oscar-Berman & Marinković, 2007; Seo et al., 2016).

Many of these same behavioral and cognitive impairments have been recapitulated in preclinical rodent models of chronic ethanol exposure. For example, heightened impulsivity (Irimia et al., 2014; Starski et al., 2020; Walker et al., 2011), deficits in behavioral flexibility (Badanich et al., 2011; Hu et al., 2015; Mejia-Toiber et al., 2014; Trantham-Davidson et al., 2014; Varodayan et al., 2018), impairments in fear extinction and retrieval (Holmes et al., 2012), and increased anxiety-like behavior (Hughes et al., 2021; Mejia-Toiber et al., 2014; Pleil et al., 2015; Starski et al., 2020; Varodayan et al., 2018) have been observed in rodents exposed to chronic ethanol. In some studies, these deficits are associated with physiological neuroadaptations in prelimbic (PL) (Holmes et al., 2012; Hughes et al., 2021; Kroener et al., 2012; Pleil et al., 2015) and infralimbic (IL) (Chuong et al., 2023; Flores-Ramirez et al., 2023; Pleil et al., 2015; Varodayan et al., 2018) subregions of the mPFC. Of note, while few studies report data from both mPFC subregions in the same experiment, those that do have found that PL and IL are differentially affected by chronic ethanol exposure (Pleil et al., 2015; Varodayan et al., 2018).

In addition to subregion-specific effects, accumulating evidence suggests that distinct mPFC cell types and circuits are also differentially affected by chronic ethanol exposure. For example, mPFC neurons expressing corticotropin-releasing factor receptor type 1 (CRF1^+^) were differentially altered by chronic ethanol exposure compared to neighboring CRFR1^-^ neurons (Patel et al., 2022) Importantly, findings from this study suggested that ethanol-induced neuroadaptations occurring in CRF1^+^ mPFC neurons likely play a role in mediating withdrawal-associated anxiety-like behavior (Patel et al., 2022). Distinct subtypes of mPFC interneurons (e.g., parvalbumin, somatostatin, VIP) are also differentially affected by chronic ethanol exposure (see Fish & Joffe, 2022 for review). In addition, recent work found that chronic ethanol exposure results in a significant increase in glutamate release onto dorsomedial PFC (i.e., PL) neurons projecting to the basolateral amygdala (McGinnis et al., 2020). In contrast, glutamate release onto basolateral amygdala-projecting ventromedial PFC (i.e., IL) neurons was significantly reduced compared to controls following the same ethanol exposure paradigm (McGinnis et al., 2020). Altogether, these data suggest that chronic ethanol exposure produces subregion- and circuit-specific neuroadaptations in PL and IL PFC.

The rostromedial tegmental nucleus (RMTg) is a small GABAergic nucleus characterized for its role in aversive signaling (Jhou, Fields, et al., 2009; Jhou, Geisler, et al., 2009; Li et al., 2019). Since its discovery in 2009, RMTg activity has been implicated in signaling the aversive properties of ethanol (Glover et al., 2016) as well as the aversive state associated with alcohol withdrawal. For example, c-Fos expression is enhanced in RMTg neurons during acute withdrawal from chronic ethanol exposure (Fu et al., 2019; Glover et al., 2019). Additionally, pharmacological or chemogenetic inactivation of the RMTg reduces anhedonia-, depression-, and anxiety-like behaviors observed after chronic ethanol exposure (Fu et al., 2019; Glover et al., 2019). Together, these data suggest that ethanol-induced changes in RMTg activity regulate symptoms of withdrawal. Importantly, recent work revealed the presence of dense input from layer 5 PL and IL mPFC glutamatergic neurons to the RMTg (Glover et al., 2023). While exposure to an aversive nociceptive stimulus produces physiological and morphological changes in RMTg-projecting PL mPFC neurons that are consistent with a loss of top-down modulation of RMTg activity (Glover et al., 2023), the effect of chronic ethanol exposure on this circuitry is unknown. The current study was designed to fill this gap by investigating the physiological neuroadaptations resulting from chronic ethanol exposure and withdrawal in distinct RMTg-projecting mPFC circuits. Our findings reveal differential effects of chronic ethanol exposure on RMTg-projecting mPFC neurons arising from PL and IL subregions. By systematically examining the effects of chronic ethanol exposure on both RMTg-projecting and projection-undefined layer 5 neuronal populations, we further reveal the circuit specificity of the long-lasting neurophysiological adaptations observed in RMTg-projecting mPFC neurons during protracted withdrawal.

## Materials & Methods

### Animals

Male Long-Evans rats (Envigo, Indianapolis, IN) were P60 upon arrival and were allowed to acclimate for at least one week prior to experimental procedures. All rats were singly housed in standard cages in a temperature-controlled room on a reverse 12:12 h reverse light/dark cycle (lights on at 22:00). Food (Teklad 7912, Envigo) and water were provided *ad libitum* for the duration of the experiment. All experimental procedures were approved by the University of Illinois Chicago Institutional Animal Care and Use Committee and adhered to the NIH Guidelines for the Care and Use of Laboratory Animals.

### Fluorescent labeling of RMTg-projecting neurons

For stereotaxic surgery, rats were induced with isoflurane (5%) and then maintained under a surgical plane of anesthesia (2-4%) for the duration of the surgery. A craniotomy was made above the right RMTg and a unilateral intracranial injection of 800 nL green fluorescent retrobeads (Lumafluor, Durham NC, USA) was made into the RMTg using coordinates corresponding to AP:-7.0; ML:+1.2; DV:-8.4 according to Paxinos & Watson (2007) at a 6° lateral angle and a rate of 10 nL/s using a custom glass pipette connected to a nanojector (Drummond Scientific Company, Broomall, PA, USA). Rats were monitored post-operatively for two weeks following surgery.

### Chronic intermittent ethanol exposure & blood ethanol concentration measurement

After at least 7 days of recovery from surgery, rats were rendered ethanol dependent using a standard chronic intermittent ethanol (CIE) exposure paradigm as previously described (Glover et al., 2019, 2021; Ramirez et al., 2024). Rats received either vaporized ethanol (CIE group) or room air (AIR controls) for 14 h/d (16:00-8:00) for 14 consecutive days. Behavioral signs of intoxication were assessed daily in CIE-exposed rats using a previously-published subjective rating scale (Glover et al., 2019, 2021; Ramirez et al., 2024) ranging from 1 (no signs of intoxication) to 5 (total loss of consciousness). Blood samples (40 μL) were collected from CIE-exposed rats via tail nick four times during the 14-day exposure period. No blood samples were collected from AIR-exposed rats; however, a tail pinch was administered to AIR controls to mimic the temporary discomfort of the tail nick experienced by CIE-exposed rats. Immediately following blood collection, samples were centrifuged (10,000 x g, 10 min, 4°C). Plasma supernatant (20 μL) was aliquoted into 0.5 mL microcentrifuge tubes and stored at −20°C until ready for processing. Blood ethanol concentration (BEC) was measured using an Analox Alcohol Analyzer (Analox Instruments Ltd, United Kingdom) within 30 days of sample collection.

### Whole-cell patch-clamp slice electrophysiology

Rats were returned to the colony after their final vapor exposure session and left undisturbed for 7-10 days, at which point they were euthanized for whole-cell patch-clamp slice electrophysiology recordings. Rats were anesthetized with isoflurane and rapidly decapitated. Brains were removed and immediately placed in cold artificial cerebrospinal fluid (ACSF) cutting solution containing (mM): NaCl (125), KCl (2.5), NaH_2_PO_4_ (1.25), NaHCO_3_ (25), glucose (10), ascorbic acid (0.4), MgCl_2_ (0.008), and CaCl_2_ (0.002). Coronal slices containing the mPFC (300 μm) were made using a vibratome (Leica Microsystems) and incubated at 34°C in continuously-oxygenated ACSF (95% oxygen/5% carbon dioxide) for at least 1 h prior to experiments. Following incubation, slices were transferred to a submerged recording chamber, held at 34°C, and perfused at a rate of 2 mL/min with oxygenated recording ASCF solution containing (mM): NaCl (125), KCl (2.5), NaHCO_3_ (25), glucose (10), ascorbic acid (0.4), magnesium chloride (0.003), and calcium chloride (0.004). The pH of all ACSF was adjusted to 7.2-7.4 and osmolarity was measured to be ∼300 mOsm.

Recordings were made with an Axon Multiclamp 700B amplifier, digitized at a sampling rate of 10 kHz with either an Instrutech LIH 8+8 (HEKA Instruments) controlled by AxographX software (Axograph Scientific) or an Axon Digidata 1550A (Molecular Devices) digitizer controlled by pClamp 11 software (Molecular Devices) running on Windows 10. Neurons were visualized using infrared interference-contrast microscopy (Olympus America), and recordings were obtained from layer 5 pyramidal neurons in the PL or IL subregions of the mPFC. RMTg-projecting neurons were identified using a GFP filter to visualize green retrobead-expressing neurons. Projection-undefined neurons were identified as GFP-cells with morphology and membrane properties characteristic of layer 5 pyramidal neurons (van Aerde & Feldmeyer, 2015). Patch pipettes were constructed from thin-walled borosilicate capillary glass tubing (Warner Instruments) and pulled with a P-97 pipette puller (Sutter Instrument Co.) to a tip resistance of 3-5 MΩ. For all recordings, neurons were left to dialyze for 15 minutes after break-in before starting experiments. Membrane properties were assessed at break-in, after 15 minutes, and at the end of each experiment. Any recordings for which the access resistance changed by greater than 20% over the course of the experiment were discarded.

For current clamp experiments, patch pipettes were filled with a potassium gluconate intracellular solution containing (mM): K-gluconate (125), KCl (20) HEPES (10), ethylene glycol tetraacetic acid (EGTA, 1), magnesium adenosine triphosphate (Mg-ATP, 2), sodium guanosine triphosphate (Na-GTP, 0.3), phosphocreatine (10), and magnesium chloride (0.02), with pH adjusted to 7.3 and osmolarity measured at ∼285 mOsm. To analyze firing characteristics and intrinsic excitability, current steps were applied for 500 ms and increased in 20 pA increments from −20 pA to 300 pA while holding the membrane potential at −70 mV. Action potential firing and intrinsic excitability data were analyzed offline using Easy Electrophysiology software.

For voltage clamp experiments, patch pipettes were filled with a cesium methanesulfonate intracellular solution containing (mM): cesium methanesulfonate (135), KCl (20), magnesium chloride hexahydrate (1), EGTA (0.2) Mg-ATP (4), Na-GTP (0.3), phosphocreatine (20), and QX-314 (2.91) with pH adjusted to 7.3 and osmolarity measured at ∼290 mOsm. To record spontaneous excitatory and inhibitory synaptic activity within the same cells, neurons were clamped at −55 mV for five minutes to record spontaneous excitatory postsynaptic currents (sEPSCs), and then clamped at +10 mV for five minutes to record spontaneous inhibitory postsynaptic currents (sIPSCs). Synaptic event frequency and amplitude were analyzed using Axograph software and Minianalysis software (SynaptoSoft Inc.). To determine the relative balance of excitatory (E) and inhibitory (I) neurotransmission, two methods were used. E/I frequency ratio (sEPSC frequency/sIPSC frequency) within each cell was used to identify the balance of excitatory and inhibitory synaptic inputs. Synaptic drive analysis [(sEPSC frequency*amplitude)/(sIPSC frequency*amplitude)] was used to determine the overall balance of excitatory and inhibitory neurotransmission within each cell.

### Histology

During acute slice preparation, the caudal portion of the brain containing the RMTg was blocked off and immersion fixed overnight in 4% paraformaldehyde. Brains were frozen on dry ice and stored at −80°C until ready for processing. Brains were sliced at 40 μm using a cryostat (ThermoScientific) and slices were mounted onto plus-charged slides and coverslipped with Vectashield mounting media with DAPI (Vector Laboratories). Visual inspection of RMTg slices was performed on an AxioImager.M2 microscope (Zeiss Microscopy) at 10X magnification to verify accurate targeting of the RMTg. Data from missed targets were excluded from analyses.

### Statistical Analysis

All statistical analyses performed in this study, including between-subjects and repeated-measures ANOVAs, were conducted using Prism 9 (GraphPad), with the significance threshold set at p<0.05. Following significant main effects or interactions, post hoc tests were performed as described below to detect significant differences between groups. Cells for which any given measure was greater than two standard deviations from the group mean were removed from all analyses. Sample sizes indicate the number of individual cells, but no more than 2 cells within an experimental group were recorded from the same rat. All data are presented as mean ± SEM, and individual data points are displayed except for action potential firing curves.

## Results

### CIE exposure produces pharmacologically relevant levels of intoxication

Similar to previously published work (Glover et al., 2019, 2021; Ramirez et al., 2024), CIE-exposed rats achieved an average intoxication score of 2.13 ± 0.04 and an average BEC (in mg%) of 214.30 ± 16.85. A Pearson correlation found a significant positive relationship between intoxication score and BEC (*r*=0.448, *p*=0.011) consistent with our previous work (Glover et al., 2021; Ramirez et al., 2024).

### CIE exposure increases excitatory drive onto PL mPFC neurons in a circuit-specific manner

To determine the lasting effect of CIE exposure on synaptic neurotransmission in layer 5 PL mPFC neurons, we conducted a synaptic drive experiment designed to measure excitatory and inhibitory neurotransmission in the same neurons. A two-way ANOVA of sEPSC frequency found significant main effects of cell population [*F(1, 30) = 6.89*, *p* = 0.014] and of vapor exposure [*F*(1, 30) = 4.24, *p* = 0.048] as well as a population by exposure interaction [*F*(1, 30) = 10.01, *p* = 0.004]. *Post hoc* comparisons with Bonferroni correction revealed that the effect of CIE exposure was specific to RMTg-projecting PL mPFC neurons, with a significant increase in sEPSC frequency observed in this population (*p* = 0.005) but not in projection-undefined neurons (*p* > 0.05, Figure 2A). No between-group differences were observed in sEPSC amplitude using the same method of analysis (*p* > 0.05, Figure 2B). CIE exposure also had no lasting effect on sIPSC frequency or amplitude in either cell population (Figure 2B-D, all *p* values > 0.05). Although CIE exposure increased sEPSC frequency in RMTg-projecting PL mPFC neurons, two-way ANOVAs did not find a significant between-group difference in presynaptic E/I balance or overall synaptic drive (*p >* 0.05, Figure 2E-F). Together, these data suggest that CIE exposure increases presynaptic glutamate release onto RMTg-projecting PL mPFC neurons, further suggesting that this cortico-subcortical circuit may be vulnerable to the lasting effects of CIE exposure.

**Figure 1.**
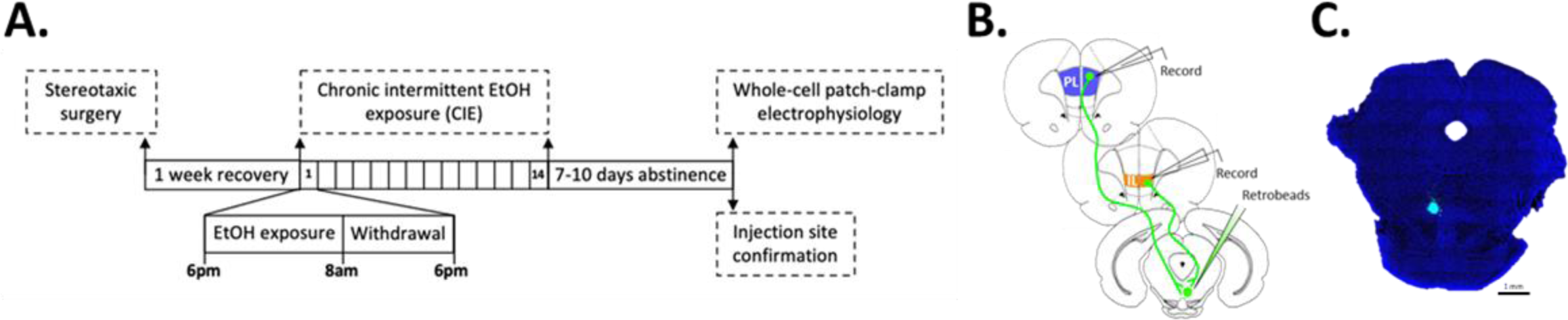
Experimental design. **(A)** Rats underwent stereotaxic surgery to inject fluorescent retrobeads into the RMTg followed by CIE exposure one week after recovery. Whole-cell patch-clamp recordings were made 7-10 days after the final vapor exposure session, followed by histological confirmation of RMTg targeting. **(B)** Recordings were made from prelimbic (blue) and infralimbic (orange) subregions of the mPFC. **(C)** Representative DAPI (blue) labeled slice showing retrobead injection site (green) in the RMTg. Scale bar= 1 mm.

**Figure 2.**
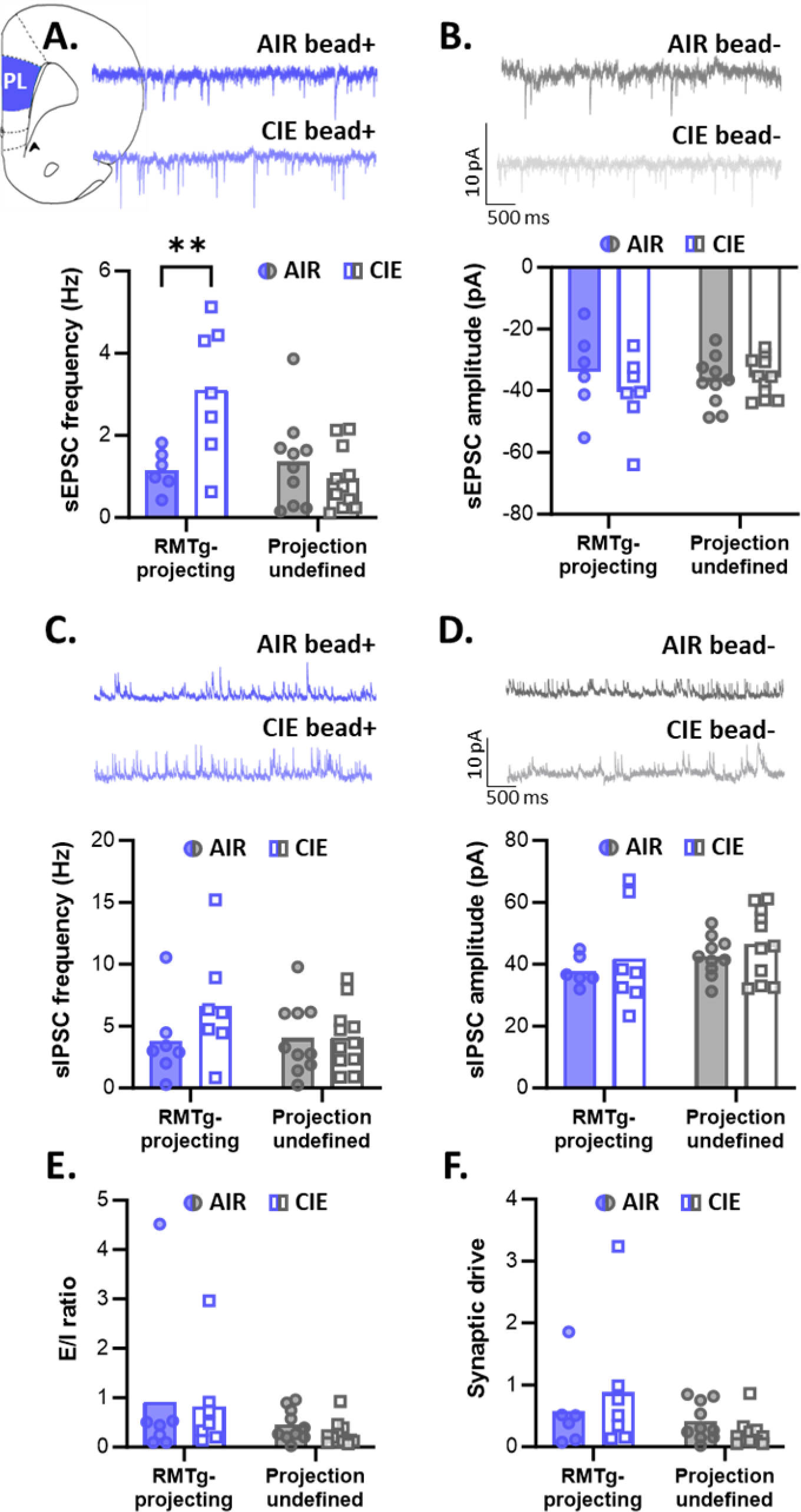
CIE-induced synaptic neuroadaptations in RMTg-projecting and projection-undefined PL mPFC neurons. **(A)** CIE exposure increased the frequency of sEPSCs, but only in RMTg-projecting PL mPFC neurons. **(B)** CIE exposure did not affect sEPSC amplitude. Representative sEPSCs are depicted above A and B. **(C)** The frequency and **(D)** amplitude of sIPSCs were not affected by CIE exposure. Representative sIPSCs are depicted above C and D. CIE also had no effect on **(E)** excitatory/inhibitory ratio and **(F)** synaptic drive. **p<0.01.

### Lasting effects of CIE exposure are subregion- and circuit-specific in the IL mPFC

In contrast to the circuit-specific effect of CIE exposure on sEPSC frequency observed in the PL mPFC, a two-way ANOVA of sEPSC frequency in IL mPFC neurons revealed a significant main effect of cell population [*F*(1, 32) = 10.53, *p* = 0.003; Figure 3A] but no effect of exposure or a population by exposure interaction (all *p* values > 0.05). Furthermore, while no significant effect of cell population or population by exposure interaction were observed with respect to sEPSC amplitude (all *p* values > 0.05), we did observe a significant main effect of vapor exposure on this measure [*F*(1, 32) = 8.59, *p* = 0.006], with greater sEPSC amplitude observed in CIE-exposed rats compared to AIR controls independent of circuit specificity.

**Figure 3.**
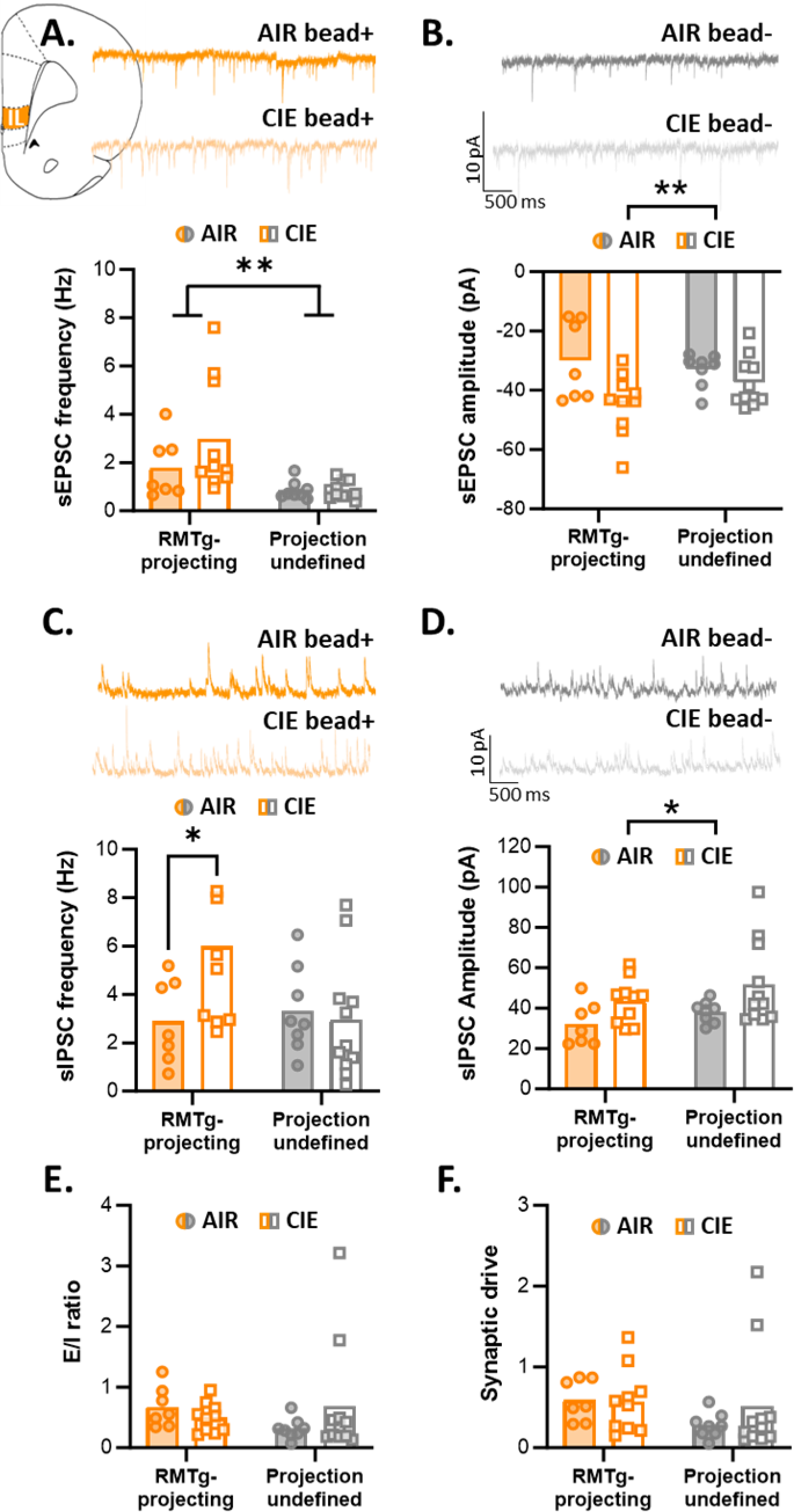
CIE-induced synaptic neuroadaptations in RMTg-projecting and projection­ undefined IL mPFC neurons. **(A)** The frequency of sEPSCs is significantly greater in RMTg­ projecting than projection-undefined IL mPFC neurons independent of vapor exposure group. **(B)** CIE exposure significantly increased sEPSC amplitude in both IL mPFC populations. Representative sEPSCs are depicted above A and B. **(C)** CIE exposure significantly increased slPSC frequency in a manner that was specific to RMTg-projecting IL mPFC neurons. **(D)** CIE exposure produced a significant increase in slPSC amplitude in both IL mPFC populations. Representative slPSCs are depicted above C and D. CIE did not affect **(E)** excitatory/inhibitory balance or **(F)** synaptic drive. *p<0.05; **p<0.01.

When looking at inhibitory neurotransmission, a two-way ANOVA of sIPSC frequency revealed a significant population by vapor exposure interaction [*F*(1, 32) =4.26, *p* = 0.047] in the absence of main effects of population or vapor exposure (all *p* values > 0.05). *Post hoc* comparisons with Bonferroni correction of the significant interaction found that CIE exposure significantly increased sEPSC frequency in RMTg-projecting IL mPFC neurons (*p* = 0.033), with no difference in projection-undefined IL mPFC neurons (*p* > 0.05; Figure 3C). Similar to our observations with respect to sEPSC amplitude, analysis of sIPSC amplitude found a significant main effect of vapor exposure [*F*(1, 32) = 6.85, *p* = 0.013] but no effect of cell population or a population by exposure interaction (all *p* values > 0.05; Figure 3D). Nevertheless, two-way ANOVAs did not uncover significant effects of cell population or vapor exposure, nor population by exposure interaction on presynaptic E/I ratio (*p* > 0.05; Figure 3E) or overall synaptic drive (*p* >0.05; Figure 3F).

Altogether, these data reveal relatively widespread effects of CIE exposure on layer 5 IL mPFC neurons, indicating postsynaptic neuroadaptations that function to increase both excitatory and inhibitory neurotransmission. This is in contrast to the relatively few effects of CIE exposure in the PL mPFC. These data further suggest that CIE exposure produces a lasting increase in inhibitory input that is specific to RMTg-projecting IL mPFC neurons.

### RMTg-projecting PL mPFC neurons have distinct intrinsic properties that are unaffected by CIE exposure

To determine whether our observed changes in synaptic strength were also accompanied by alterations in intrinsic properties, we next performed a series of current clamp recordings in layer 5 mPFC neurons from AIR- and CIE-exposed rats. A three-way ANOVA of action potential firing between vapor exposure groups and cell populations across a series of current steps found a significant main effect of cell population [*F*(1, 464) = 54.26, *p* < 0.0001] with RMTg-projecting PL mPFC neurons having reduced intrinsic excitability relative to projection-undefined PL mPFC neurons. A significant population by vapor exposure interaction was also evident [*F*(1, 454) = 5.57, *p* = 0.019] (Figure 4A), indicating that RMTg-projecting and projection-undefined PL mPFC neurons are differentially affected by CIE exposure. Based on this, we next performed separate two-way ANOVAs comparing the effect of vapor exposure on cell firing across current steps within each cell population. This analysis revealed a significant main effect of exposure in projection-undefined layer 5 PL mPFC neurons [*F*(1, 224) = 5.82, *p* = 0.017] but not in RMTg-projecting PL mPFC neurons (p > 0.05). No significant interaction was observed between vapor exposure and current step in projection-undefined PL mPFC neurons (*p* > 0.05). In addition, no significant effects of vapor exposure or interaction with current step were observed in RMTg-projecting PL mPFC neurons (all *p* values > 0.05). These data indicate that CIE exposure significantly decreased the intrinsic excitability of projection-undefined, layer 5 pyramidal cells in the PL mPFC without affecting the excitability of RMTg-projecting PL mPFC neurons

**Figure 4.**
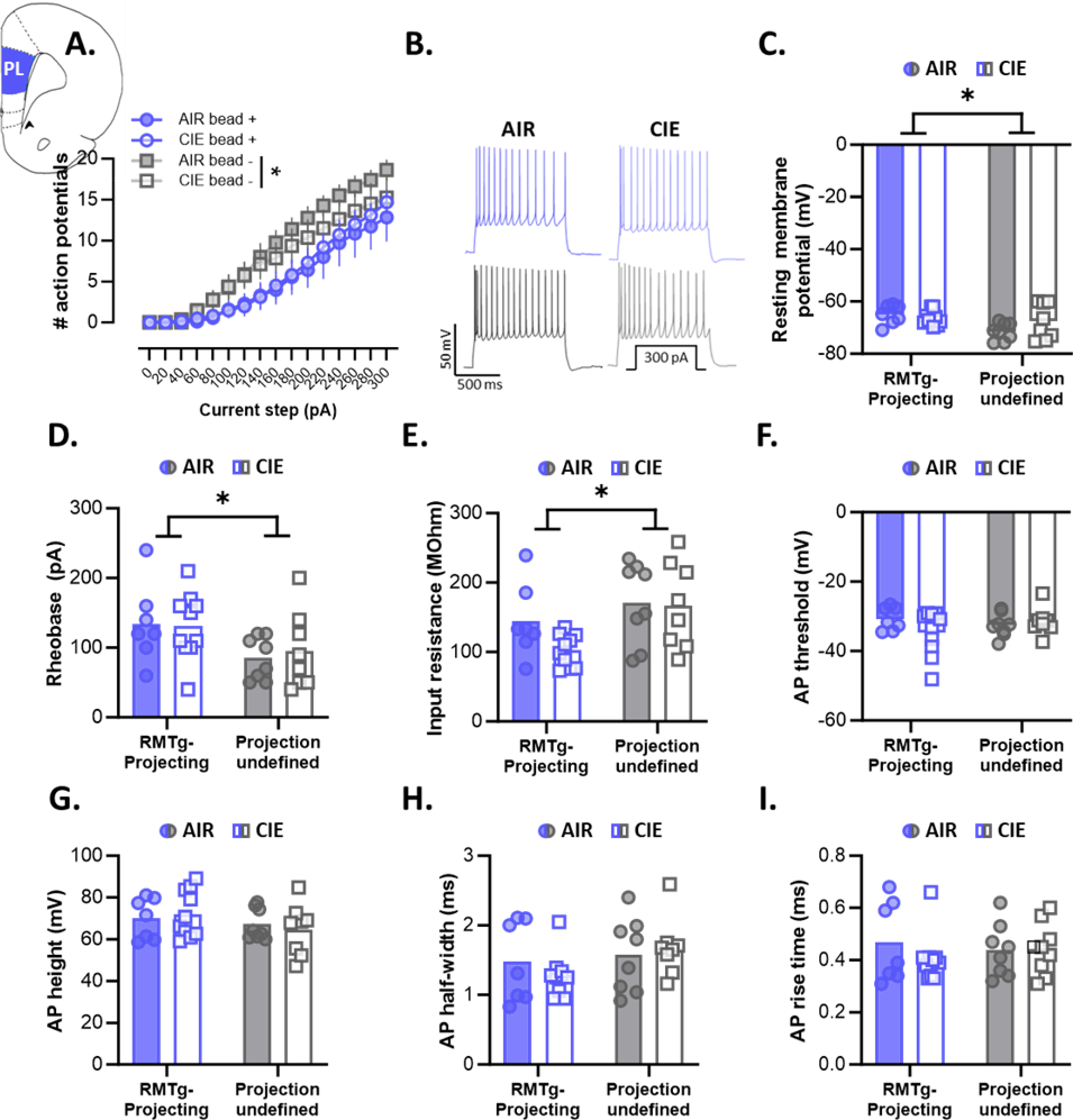
Effect of CIE exposure on intrinsic properties of RMTg-projecting and projection­ undefined PL mPFC neurons. **(A)** CIE exposure significantly decreased the number of action potentials fired across a series of current steps in projection-undefined, but not RMTg-projecting, PL mPFC neurons. **(B)** Representative traces of action potential firing at 300 pA current. RMTg­ projecting PL mPFC neurons exhibited significantly lower resting membrane potential **(C),** higher rheobase **(D),** and lower input resistance **(E)** compared to projection-undefined PL mPFC neurons, regardless of vapor exposure. There were no effects of CIE exposure on action potential firing threshold **(F),** height **(G),** half-width **(H),** or rise time (I). *p<0.05.

In contrast, CIE exposure did not have a significant lasting effect on other intrinsic properties in either RMTg-projecting or projection-undefined PL mPFC neurons. Instead, our analysis revealed a number of differences between cell populations. For example, a two-way ANOVA of resting membrane potential (RMP) found a significant main effect of cell population [*F*(1, 29) = 6.80, *p* = 0.014], with RMTg-projecting PL mPFC neurons exhibiting lower RMP than projection-undefined neurons. This occurred in the absence of an effect of vapor exposure or population by exposure interaction (all p values > 0.05; Figure 4C). Similarly, rheobase differed significantly between cell populations with significantly more current required to induce an action potential in RMTg-projecting than projection-undefined PL mPFC neurons [*F*(1, 29) = 6.31, *p* = 0.018] (Figure 4D). This was accompanied by a significant difference in input resistance between cell populations with projection-undefined PL mPFC neurons having greater resistance than RMTg-projecting PL mPFC neurons [*F*(1, 29) = 6.86, *p* = 0.014] (Figure 4E). No significant effects of vapor exposure or cell population by exposure interactions were observed in any of these measures (all *p* values > 0.05).

Finally, to determine whether CIE exposure affected discrete action potential characteristics, we performed two-way ANOVAs examining the effect of vapor exposure on action potential threshold, height, half-width and rise time. No significant main effects of vapor exposure or cell population and no exposure by population interactions were observed for any of these variables (all *p* values > 0.05; Figure 4F-I). Altogether, these findings suggest that CIE exposure produces a lasting decrease in intrinsic excitability of projection-undefined layer 5 PL mPFC neurons that is absent in RMTg-projecting PL mPFC neurons. In addition, our findings suggest that RMTg-projecting PL mPFC neurons may be a physiologically-unique population of neurons given that these neurons exhibit measures of significantly lower intrinsic excitability compared to surrounding layer 5 pyramidal cells.

### CIE exposure increases the excitability of RMTg-projecting IL mPFC neurons

Similar to our observation in PL mPFC neurons, a three-way ANOVA of action potential firing in IL mPFC neurons revealed a significant main effect of cell population [*F*(1, 464) = 68.43, *p* < 0.0001] and a significant population by vapor exposure interaction [*F*(1, 464) = 32.20, *p* < 0.0001]. Therefore, two-way ANOVAs were used to examine the differential effects of vapor exposure on cell firing across current steps within each cell population. In contrast to our observations in the PL mPFC, this analysis did not uncover a main effect of vapor exposure or interaction with current step in projection-undefined IL mPFC neurons (all *p* values > 0.05). In RMTg-projecting IL mPFC neurons, however, there was a significant main effect of vapor exposure (*F*(1, 256) = 54.18, *p* < 0.0001] and a significant vapor exposure by current step interaction (*F*(15, 256) = 3.71, *p* < 0.0001], revealing that CIE exposure significantly increased action potential firing in this neural circuit (Figure 5A). When looking at other measures of intrinsic excitability in IL mPFC neurons using two-way ANOVAs, we again found main effects of cell population, with significantly lower resting membrane potential in RMTg-projecting versus projection-undefined IL mPFC neurons [*F*(1, 29) = 4.40, *p* = 0.045] (Figure 5C). This was accompanied by a significantly higher rheobase [*F*(1, 29) = 6.04, *p* = 0.02] (Figure 5D) and a significantly lower input resistance [*F*(1, 29) = 9.43, *p* = 0.005] (Figure 5E) in RMTg-projecting IL mPFC neurons compared to surrounding layer 5 pyramidal cells. Thus, although CIE exposure significantly increased intrinsic excitability in RMTg-projecting IL mPFC neurons relative to AIR controls, these results indicate that this neural circuit has lower overall excitability compared to other IL mPFC layer 5 pyramidal cells independent of CIE exposure.

**Figure 5.**
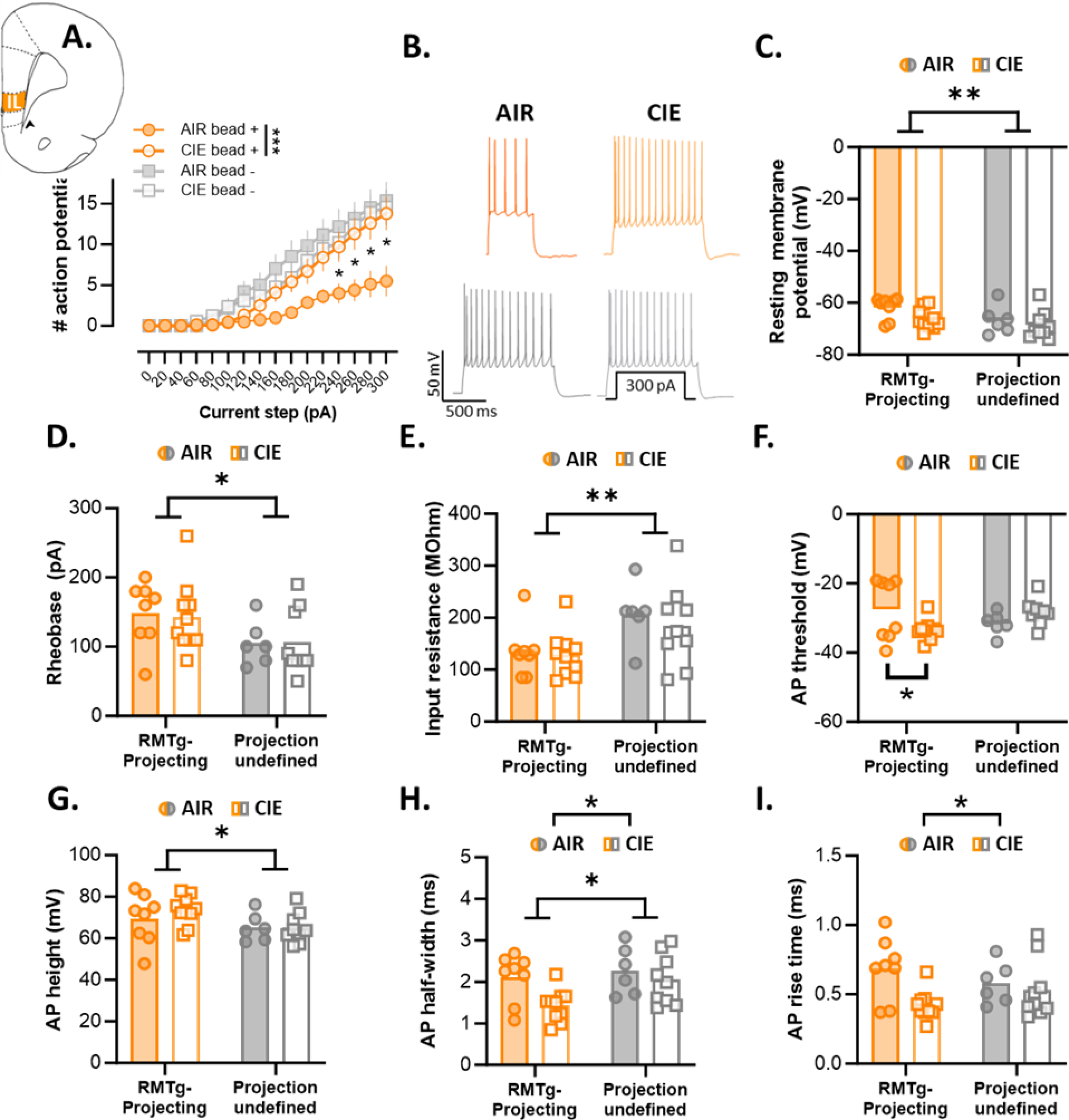
Effect of CIE exposure on intrinsic properties of RMTg-projecting and projection­ undefined IL mPFC neurons. **(A)** CIE exposure significantly increased action potential firing in RMTg-projecting, but not projection-undefined, IL **mPFC** neurons. **(B)** Representative traces of action potential firing at 300 pA current. RMTg-projecting IL mPFC neurons had significantly lower resting membrane potential **(C),** higher rheobase **(D),** and lower input resistance **(E)** compared to projection-undefined IL mPFC neurons. **(F)** CIE exposure increased the action potential firing threshold in RMTg-projecting IL mPFC neurons without affecting projection-undefined neurons. Action potential height **(G)** was significantly higher and half-width **(H)** was significantly lower in RMTg-projecting IL mPFC neurons compared to projection-undefined neurons. In addition, CIE exposure significantly deceased action potential half-width **(H)** and rise time (I) independent of cell population. *p<0.05, **p<0.01.

Examination of action potential characteristics from these cell populations revealed additional effects of CIE exposure on IL mPFC neurons. A two-way ANOVA of action potential threshold revealed a significant vapor exposure by cell population interaction [*F*(1, 29) = 6.34, *p* = 0.018]. *Post hoc* comparisons using a Bonferroni correction found that CIE exposure significantly decreased the firing threshold in RMTg-projecting (*p* = 0.041) but not projection-undefined (*p* > 0.05) IL mPFC neurons (Figure 5F), supporting our finding of a CIE-induced increase in action potential firing in the RMTg-projecting population. A two-way ANOVA for action potential height found a significant main effect of cell population [*F*(1, 29) = 4.54, *p* = 0.042], but no significant effect of vapor exposure or exposure by cell population interaction (all *p* values > 0.05; Figure 5G). Significant main effects of cell population [*F*(1, 29) = 4.20, *p* = 0.049] and of vapor exposure [*F*1, 29) = 5.65, *p* = 0.024] were also observed for action potential half-width in the absence of a significant interaction between these factors (*p* > 0.05; Figure 5H). Finally, analysis of action potential rise time also revealed a significant main effect of vapor exposure [*F*(1, 29) = 6.14, *p* = 0.019] in the absence of significant effects of cell population or a vapor exposure by population interaction (all *p* values > 0.05; Figure 5I). Taken together, these findings show that, similar to RMTg-projecting PL mPFC neurons, RMTg-projecting IL mPFC neurons have reduced intrinsic excitability compared to surrounding layer 5 pyramidal cells. However, in contrast to PL mPFC neurons, CIE exposure has a number of long-lasting effects on both RMTg-projecting and projection-undefined IL mPFC neurons.

## Discussion

Our recent work identified a dense population of layer 5 glutamatergic neurons projecting from the mPFC to the RMTg (Glover et al., 2023). This neural circuit is responsible for driving avoidance and is activated in response to aversive stimuli (Glover et al., 2023). Aversion plays a key role in critical aspects of the addiction cycle, including the negative affective state that occurs during withdrawal from chronic alcohol exposure (Koob & Volkow, 2016). Findings from the current study show that exposure to chronic ethanol produces differential long-lasting changes in the synaptic physiology and intrinsic excitability of RMTg-projecting PL and IL mPFC projections, providing support for the hypothesis that alterations in these circuits play a key role in aversive signaling during withdrawal from chronic ethanol exposure.

In the present study, we found that CIE exposure selectively increased sEPSC frequency in RMTg-projecting PL mPFC neurons. In addition, decreased intrinsic excitability was observed in population-undefined PL mPFC neurons after CIE exposure, whereas intrinsic measures were unaffected in RMTg-projecting PL neurons. These results differ from previous work, which found no change in sE/IPSCs or intrinsic excitability 7 days into withdrawal from chronic ethanol exposure (Trantham-Davidson et al., 2014). However, our findings are in partial agreement with findings from Galaj et al. (2020), which also observed a decrease in the intrinsic excitability of PL mPFC neurons 21 days after chronic ethanol exposure. This was accompanied by an increase in both the frequency and amplitude of sEPSCs (Galaj et al., 2020). Although the effect of CIE exposure on sEPSCs in the PL mPFC was limited to a change in frequency in RMTg-projecting PL neurons, it should be noted that a nonsignificant increase in sEPSC amplitude was also observed in this same cell population in CIE-exposed rats relative to AIR controls. It is unclear what precisely contributes to the discrepant findings between the present study and previous work. However, neither Galaj et al. (2020) nor Trantham-Davidson et al. (2014) characterized neuronal populations by their projection targets. This may be particularly important given the circuit-specific findings from the current study in combination with other work in male rats showing that chronic ethanol exposure increased presynaptic excitatory regulation that was specific to “Type A” layer 5 PL neurons (Hughes et al., 2021), which are subcortically projecting neurons and distinct from “Type B”, cortico-cortical layer 5 projection neurons (Lee et al., 2014).

In the IL mPFC, withdrawal from chronic ethanol exposure produced both circuit-specific and -nonspecific effects. The amplitude of sEPSCs and sIPSCs was significantly increased in CIE-exposed layer 5 IL mPFC neurons independent of circuit specificity. However, RMTg-projecting IL mPFC neurons also exhibited a significant increase in sIPSC frequency and in intrinsic excitability that was absent in neighboring, projection-undefined neurons. To our knowledge, only two other studies have explored the effect of chronic ethanol exposure on layer 5 pyramidal neurons in the IL mPFC, and both restricted their analyses to changes in inhibitory synaptic input. In the first study, Flores-Ramirez et al. (2023) saw no changes in sIPSC frequency or amplitude 8 hr into acute withdrawal from chronic ethanol exposure. This is in contrast to Chuong et al. (2023), which observed a significant increase in both the frequency and amplitude of sIPSCs after CIE exposure. Although seemingly in at least partial agreement with the current findings, Chuong et al. (2023) performed recordings within an hour after the final ethanol vapor exposure session and allowed slices containing the IL mPFC to undergo acute withdrawal *in vitro*. In addition, neither study distinguished between distinct cortico-subcortical projections, making comparisons between the current findings and these studies difficult. Nevertheless, taken together, the previous and current work examining the effect of chronic ethanol exposure on PL and IL mPFC physiology makes clear that duration of abstinence following chronic ethanol exposure is a critical driver of the neurophysiological adaptations observed.

While few physiological studies have included analysis of both PL and IL subregions, those that have provide strong evidence that the effects of chronic ethanol exposure are frequently subregion-(Pleil et al., 2015; Varodayan et al., 2018) and circuit-specific (McGinnis et al., 2020). The current findings lend further support to this notion by revealing the unique vulnerability of IL mPFC neurons to the long-lasting effects of withdrawal from chronic ethanol relative to PL mPFC neurons. Indeed, CIE exposure exerted significant changes in both synaptic neurotransmission and intrinsic excitability in IL mPFC neurons that were substantially more modest or entirely absent in the PL mPFC. Similarly, our data show that RMTg-projecting mPFC neurons, particularly those originating in the IL mPFC, exhibit long-lasting changes as a result of chronic ethanol exposure that are not evident in neighboring, projection-undefined populations. Mechanistically linking these distinct subregion- and circuit-specific effects with changes in mPFC-mediated behaviors will be crucial to provide insight into the precise ways in which these physiological neuroadaptations contribute to maladaptive behaviors that facilitate continued drinking.

In addition to CIE-induced neuroadaptations, we also identified significant physiological differences between RMTg-projecting and projection-undefined neuronal populations within both mPFC subregions. This included lower resting membrane potential, input resistance, and rheobase in RMTg-projecting mPFC neurons of either subregion relative to neighboring, projection-undefined neurons. Together, these measures contribute to lower overall intrinsic excitability in RMTg-projecting circuits relative to surrounding neurons. The possibility that RMTg-projecting mPFC neurons constitute a physiologically distinct subpopulation of pyramidal neurons is supported by previous studies. For example, one study looking at layer 2/3 pyramidal CRF1^+^ neurons found that these neurons exhibit a unique physiological signature compared to surrounding CRF1^-^ neurons. Interestingly, these two cell populations were also differentially affected by chronic ethanol exposure (Patel et al., 2022). It is also possible that our recordings of population-undefined mPFC neurons are comprised, at least in part, of cortico-cortical, Type-B, neurons, also referred to as thin-or slender-tufted neurons (Lee et al., 2014; van Aerde & Feldmeyer, 2015). Indeed, the intrinsic properties of RMTg-projecting PL and IL mPFC neurons reported here align well with those of Type-A, or thick-tufted, layer 5 mPFC neurons, which are the least excitable of the layer 5 mPFC neuron subtypes (van Aerde & Feldmeyer, 2015). Nevertheless, we also observed a significant difference in sEPSC frequency between RMTg-projecting and projection-undefined IL mPFC neurons suggesting the possibility that at least some mPFC neurons projecting to the RMTg are distinct with respect to both intrinsic properties and synaptic input.

The current investigation did not include data from both sexes, therefore limiting our conclusions to male rats. There is a large gap in our understanding of the structure and function of the female mPFC, and emerging evidence supports the notion of at least some degree of functional differentiation of the mPFC between males and females [for review, see (Laine et al., 2024)]. Additionally, recent work has uncovered important sex differences in ethanol-induced physiological neuroadaptations in the mPFC (Avchalumov et al., 2021; Hughes et al., 2021; Joffe et al., 2020). One potential reason for this is that female rats require higher levels of intoxication to reach a similar level of behavioral impairment compared to males (Glover et al., 2021). Because of this, preclinical models that utilize identical exposure paradigms for both sexes may not produce the same degree of behavioral or physiological deficits. Thus, future studies in that examine the effects of ethanol exposure at levels of intoxication that match the current study in females compared to a separate group of females exposed to a higher level of intoxication that produces a similar degree of behavioral impairment observed in males will be crucial for developing informed conclusions regarding the potential sex-dependent effects of chronic ethanol exposure on mPFC physiology.

In summary, the current study provides the first-ever investigation of the long-lasting effects of chronic ethanol exposure on RMTg-projecting mPFC neurons while also identifying physiological characteristics that distinguish this neural circuit from neighboring layer 5 neurons. Our data reveal distinct subregion- and circuit-specific neuroadaptations in the PL and IL mPFC 7-10 days following cessation of chronic ethanol exposure. In particular, RMTg-projecting neurons originating in the IL mPFC appear uniquely vulnerable to withdrawal-induced physiological alterations compared to both projection-undefined IL mPFC neurons and PL mPFC neurons of both populations. Altogether, these data suggest that long-lasting physiological alterations in cortico-subcortical circuits that regulate aversive signaling may contribute to maladaptive behaviors that impede recovery from chronic alcohol exposure.

## Acknowledgements

The authors thank Hyerim Yang and Nathaly Arce Soto for technical assistance. This work was supported by the following grants from the National Institute on Alcohol Abuse and Alcoholism at the National Institutes of Health: R00 AA024208 (EJG), R01 AA029130 (EJG), T32 AA026577 (KRP; JES).

